# TRAIP regulates DNA double-strand break-induced ATM activation

**DOI:** 10.1101/2021.10.11.462297

**Authors:** Tobias Gleich, Manfredo Quadroni, Gökhan Yigit, Bernd Wollnik, Marcel Huber, Christine Pich, Daniel Hohl

**Author notes:** To whom correspondence should be addressed to Daniel Hohl.

## Abstract

DNA double-strand breaks (DSBs) affect cell survival and genomic integrity. They are repaired by a highly coordinated process called the DNA damage response. Here, we report that the ubiquitously expressed nucleolar E3 ubiquitin ligase TRAF-interacting protein (TRAIP), previously shown to regulate the spindle assembly checkpoint, has an essential role during the DNA damage response. A biotinylation proximity screening assay (BioID) identified Ku80, Ku70, SMARCA5 (SNF2H) and DNA-PKcs as novel TRAIP interactors. Co-immunoprecipitations demonstrated that the interaction of TRAIP with Ku80 was transiently increased while the one with SMARCA5 was strongly decreased after treatment of HeLa cells with neocarzinostatin (NCS). Treatment of fibroblasts from a microcephalic primordial dwarfism patient carrying a hypomorphic TRAIP mutation or shRNA-mediated knockdown of TRAIP in HeLa cells with NCS impaired the activation of ataxia-telangiectasia mutated (ATM), a protein kinase crucial for the DNA damage response. As consequence, the maintenance of γH2AX and Chk2-T68 phosphorylation, two downstream targets of ATM, was significantly abrogated after NCS-inflicted DSBs. DNA repair assays showed that TRAIP inhibits incorrect end utilization during non-homologous end joining. These observations highlight TRAIP as novel regulator of ATM activity in DNA damage signaling.

## INTRODUCTION

DNA double-strand breaks (DSBs) caused by irradiation, chemical compounds or collapsing replication forks are the most toxic DNA lesions. Unrepaired DSBs potentially result in genomic instability leading to mutations, chromosome fusion, deletion or translocation (1). To ensure efficient repair of DNA lesions complex pathways evolved during evolution. DNA damage sensing and the timely and spatial recruitment of DNA repair factors have to be tightly controlled and coordinated (2). DSBs are mainly repaired by two pathways which are active at different time-points during the cell cycle. While error-free homologous repair (HR) is favored during S and G2 phase, error-prone non-homologous end-joining (NHEJ) is mainly active during G1 (3).

Upon DNA damage, chromatin remodeling and recruitment of DNA repair proteins is regulated by posttranslational modifications. Immediately after the formation of DSBs, ATM is activated (4) and phosphorylates histone H2AX on residue S139 to form γH2AX in the chromatin surrounding DNA lesions (5). This event is crucial since γH2AX is needed to recruit additional DNA repair proteins such as MDC1 or 53BP1 (6). The formation of γH2AX is important for the DNA damage repair pathway choice, the repair of the DNA lesion itself and DNA damage checkpoint signaling (7). In the initial step of DNA repair by NHEJ, DSBs are recognized with high affinity and very rapidly bound in a sequence-independent manner by the ring-shaped Ku70/Ku80 heterodimer to protect DNA from nucleolytic degradation (8). Upon binding of Ku70/Ku80 to DNA, DNA-PKcs attaches to the heterodimer resulting in the activation of the DNA-PK catalytic activity (9).

The TRAIP protein is a functional E3 ubiquitin ligase with a RING domain at the N-terminal end (10). If ectopically expressed, TRAIP interacts with the TRAF signaling complex (11). The Drosophila homologue NOPO (no poles) is essential for embryo development (12) and ablation of TRAIP in mice results in embryonic lethality caused by decreased proliferation and increased apoptosis (13). Knockdown experiments demonstrated that TRAIP is crucial for proliferation (14) and the mitotic spindle assembly checkpoint (15). TRAIP depletion leads to chromosomes defects in anaphase and decreases MAD2 levels on kinetochores (15). Examining the function of TRAIP in meiosis revealed that TRAIP colocalizes to the centromere region, affects MAD2 levels at centromeres, and negatively regulates the abundance of the MAD2 substrate securin (16).

Recently, TRAIP was found to be implicated in repair and replication of DNA. Hypomorphic mutations in *TRAIP* were identified in three families affected by microcephalic primordial dwarfism (MPD) (17). Fibroblasts from these patients demonstrated reduced phosphorylation of H2AX (γH2AX; H2AX-pS139) and RPA2 (pS2 and pS4) after UVC radiation, leading to increased replication fork stalling (17). A crucial function of TRAIP in DNA replication was confirmed in two other reports showing an interaction of TRAIP with PCNA (18,19). TRAIP binds to RNF20 and to RAP80 which is important for TRAIP translocation to DSBs and the recruitment of the BRCA1-A complex to DNA damage sites, respectively (20).

To find novel TRAIP partners we used a proximity-dependent biotinylation assay (BioID) which can identify direct and indirect, e.g. when participating in a common complex, interactions partners for a protein-of-interest (21). A large proportion of significant proteins emerging from this BioID screen are involved in DNA repair and replication. In particular, we identified the core NHEJ proteins, DNA-PKcs, Ku70, and Ku80, and the chromatin remodeler SMARCA5 as highly significant hits. We confirmed Ku80 and SMARCA5 as novel TRAIP interaction partners by coimmunoprecipitation. Neocarzinostatin-induced activation of ATM and phosphorylation of H2AX and Chk2, ATM downstream targets, were strongly reduced in fibroblasts from MPD patients with hypomorphic TRAIP mutations and in TRAIP-depleted HeLa cells, indicating an important role for TRAIP in DNA damage-mediated ATM activation.

## MATERIAL AND METHODS

### Cell culture and production of lentivirus

HeLa, 293T, and 293FT (Invitrogen) cells and primary human fibroblasts were cultured in 3T3 medium (14). Production of lentivirus was performed in 293FT cells as described (14).

### Plasmids

The following plasmids were used: pcDNA3.1 MCS-BirA_R118G_-HA (Addgene #36047, gift from Dr. K. Roux, (22)), pCBASceI (Addgene #26477, gift from Dr. M. Jasin, (23)), pimEJ5GFP (Addgene #44026, gift from Dr. J. Stark, (24)), pDRGFP (Addgene #26475, gift from Dr. M.Jasin, (25)), pEGFP-C1-FLAG-Ku80 (Addgene #46958, gift from Dr. S. Jackson, (26)), pCMV6-SMARCA5-Myc-DDK (Origene), pLVX-mCherry-N1 (Clontech), pCDH-EGFP-TRAIP (15), pCMV-HATRAIP (27). Lentiviral shRNA vectors SHC, sh742, and sh743 (MISSION, Sigma) were described previously as Control B, T4 and T5, respectively (14). TRAIP full-length coding sequence without the termination codon was amplified from pmycTRAIP (27) and cloned into the AgeI/BamHI sites of pcDNA3.1 MCS-BirA_R118G_-HA. The product was verified by sequencing.

### γH2AX foci kinetics

HeLa cells were seeded into 12-well plates and transduced with lentivirus SHC2, sh742 or sh743. Puromycin (2μg/ml) selection started 24 hours later. Cells were treated with neocarzinostatin (200ng/ml) 48 hours post-transduction, fixed at the indicated time-points and processed for immunfluorescence staining as described below. The number of γH2AX foci was counted using ImageJ.

### Cell fractionation and immunoblot analysis

Whole cell lysate were prepared in 1% SDS/phosphate buffered saline, sonication (2 × 10 sec), heating for 5 min at 85°C and centrifugation for 20 min at 13000rpm at 4°C. Supernatants were analyzed by immunoblot using the following antibodies: mouse anti-γH2AX (05-636, Millipore) 1:1000, rabbit pS1981-ATM (ab81292, Abcam) 1:1000, rabbit ATM (PC116, Millipore) 1:1000, rabbit pT68-Chk2 (2661, Cell Signaling Technology) 1:3000, rabbit-anti TRAIP (151307, Abcam) 1:5000, goat anti-TRAIP (3193, Imgenex) 1:2000, mouse anti-FLAG (F1804, Sigma) 1:2000, rabbit anti-HA (sc-805, Santa Cruz Biotechnologies) 1:1000, mouse anti-nucleolin C23 (sc-8031, Santa Cruz Biotechnology) 1:1000, mouse anti-α-tubulin (T9026, Sigma) 1:1000, rabbit anti-actin (A2066, Sigma) 1:2000. Cell fractionation was carried out using 2×10^6^ HeLa cells (28).

### Immunofluorescence analysis

HeLa cells were grown on glass cover-slips and were, if not otherwise indicated, fixed for 20 min in 4% paraformaldehyde/Tris-buffered saline (TBS) and permeabilized with 0.2% Triton-X100/TBS for 10 min. Subsequently, cells were blocked in 10% normal horse or normal goat serum, incubated for 2 hours with primary and 1 hour with secondary antibodies and 1 min in DAPI (0.5μg/ml). The following antibodies were used: mouse anti-γH2AX (Millipore) 1:1000 and Alexa594 goat anti-mouse (Invitrogen) 1:600. Images were acquired using a LSM700 confocal laser scanning microscope (Zeiss) with laser diodes 405/488/555, SP490/SP555/LP560 emission filters, 2 PMT detectors through a 63x/1.40 oil objective and analyzed by Zen2010 software (Zeiss).

### Laser microirradiation

HeLa cells expressing TRAIP-GFP were cultured for 24 hours in the described medium supplemented with 10μM 5-bromo-2’-deoxyuridine. The selected cell was zoomed in 5-fold and the ROI was exposed to 100% output of a 405nm laser with a scan speed of 25.21μs/pixel. Cells were recorded 30sec before laser-microirradiation and every 30 sec afterwards through an Apochromat 63x Water-Oil immersion objective with a 1.2 NA on a confocal LSM 700 (Zeiss) with a CoolSNAPHQ2 camera and analyzed by the Efficient Navigation software (Zeiss).

### Auto-ubiquitination assay

293T cells were seeded in six well plates (6×10^5^ cells/well) and transfected the next day with 1μg FLAG-Ubiquitin and 1μg HA-TRAIP or TRAIP-BirA*-HA as described (29). Cell extracts were prepared 48 hours later and immunoprecipitations were performed (27) using rabbit anti-HA antibody (sc-805, Santa Cruz Biotechnology). Immunoprecipitates were analyzed by immunoblot using mouse anti-Flag M2 antibody (F1804; Sigma) and rabbit anti-HA antibody (sc-805; Santa Cruz Biotechnology) by chemiluminescence detection.

### RNA isolation and qRT-PCR

RNA was isolated using the RNeasy Mini Kit (Qiagen). The cDNA was synthesized using Primescript RT reagent kit (TakaRa), and qPCR analysis was carried out (14) using primers (Qiagen) for TRAIP and RPL13A (endogenous control). Amplification efficiencies (E=2±0.1) of primer pairs were calculated using 5-fold serial template dilutions and analysis of the slope by linear regression.

### Purification and mass spectrometry analysis of biotinylated proteins

27×10^6^ HeLa cells from three 10cm plates transiently transfected with TRAIP-BirA*-HA or BirA*-HA were incubated for 24 hours in medium containing 50μM biotin. Biotinylated proteins were purified using 260μl MyOne Streptavidin T1 beads (Novex Life Technologies) per 5mg total protein as described previously (22). Proteins were eluted from streptavidin beads with 40μl of 4xSDS-PAGE sample buffer at 95°C for 5 min. The eluate was separated on a 12.5% polyacrylamide gel on a distance of 2.5 cm. After fixation and Coomassie staining, entire gel lanes were excised into 5 equal regions from top to bottom and digested with trypsin (Promega) as described (30). Data-dependent LC-MS/MS analysis of extracted peptide mixtures after digestion was carried out with a LTQ-Velos PRO orbitrap mass spectrometer (ThermoFisher) interfaced to a nanocapillary HPLC (RSLC3000, Dionex) equipped with a 75μm ID, 25cm C18 reversed-phase column (Acclaim PepMap, Dionex). Peptides were separated along a 95 min gradient from 5% to 45% acetonitrile in 0.1% formic acid at 0.3 μl/min. Full MS survey scans were performed at 60’000 resolution. In data-dependent acquisition controlled by Xcalibur 2.2 software (ThermoFisher), the twenty most intense multiply charged precursor ions detected in the full MS survey were selected for collision-induced dissociation and analysis in the linear trap with an isolation window of 4.0 m/z and then dynamically excluded from further selection during 25s. Collections of tandem mass spectra for database searching were generated from raw data and searched using Mascot (Matrix Science, London, UK; version 2.1.0) against the release 2014_08 of the UNIPROT database, (SWISSPROT + TrEMBL, www.expasy.org) restricted to human taxonomy. The software Scaffold (Proteome Software Inc.) was used to validate MS/MS based peptide (minimum 95% probability (31) and protein (min 99% probability (32) identifications, perform dataset alignment and subtraction as well as parsimony analysis to discriminate homologous hits.

### Co-immunoprecipitation

4×10^6^ HeLa cells were transfected with 10μg plasmid DNA using jetPRIME (PolyPlus). 30 hours later, cells were extracted in 5ml lysis buffer (50mM Tris-HCl pH 7.4, 150mM NaCl, 1mM EDTA, 1% Triton X-100) for 30 min at RT. Supernatants after centrifugation (10 min, 12000xg, 4°C) were incubated with 30μl ANTI-FLAG M2 Affinity Gel (Sigma) overnight at 4°C. Beads were recovered by centrifugation for 3min at 500xg and 4°C and washed five times with 50mM Tris HCl pH 7.4, 150mM NaCl (TBS). Proteins were eluted from the beads according to the manufacturer’s protocol. Equal quantities of eluates were loaded on SDS-PAGE and analyzed by immunoblotting.

### DSB repair assays

Stable 293T cell lines with either pimEJ5GFP or pDRGFP were established by transfection and selection with puromycin (2μg/ml) for 14 days. 3×10^5^ cells/well seeded in 6-well plates were transduced with SHC, sh742 or sh743. Cells were transfected (29) 36 hours later either with pCBASceI and pLVX-mCherry-N1, or pLVX-mCherry-N1 or EGFP. Cells were collected 72 hours post-transfection, and the percentages of GFP- and mCherry-positive cells were measured by analyzing 1.5×10^5^ to 3×10^5^ cells by flow cytometry (FACSCalibur, BD). Results are expressed as ratio of GFP-positive to mCherry-positive cells. To calculate the fold-difference in repair efficiency each ratio was divided by the value from SHC transduced cells used as control.

### Statistical analyses

Statistical significance was calculated with the GraphPad Prism 6 software (GraphPad software Inc., La Jolla, USA). Statistical results are shown as *p<0.5, **p<0.01, ***p<0.001, ****p<0.0001.

## RESULTS

### Identification of novel TRAIP-interacting proteins

To search for interaction partners of TRAIP, we used a proximity-dependent biotin labeling system (Bio-ID). TRAIP was fused to the N-terminal end of the promiscuous prokaryotic biotin protein ligase BirA_R118G_ (BirA*) tagged with HA (TRAIP-BirA*-HA). The BirA* part of the fusion protein provided a biotinylase activity allowing to mark proteins interacting either directly with TRAIP or found in its proximity (22).

First, we investigated whether TRAIP-BirA*-HA was functionally similar to TRAIP. Ubiquitination experiments showed that TRAIP-BirA*-HA was autoubiquitinated similarly as HA-TRAIP (Suppl. Fig. 1A), indicating that the E3 ligase ubiquitin of TRAIP-BirA*-HA was functional. Analysis of HeLa cells transiently transfected with TRAIP-BirA*-HA revealed the presence of the expected 80kDa TRAIP-BirA*-HA fusion protein (Suppl. Fig. 1B). Streptavidin-HRP immunoblot analysis of transfected cells cultured for 24 hours in biotin supplied medium detected specific signals of biotinylated proteins in TRAIP-BirA*-HA cells which were different from BirA*-HA transfected cells (Suppl. Fig. 1B). Confocal immunofluorescence microscopy of TRAIP-BirA*-HA transfected HeLa cells using anti-HA or anti-TRAIP antibodies showed that the TRAIP-BirA* fusion protein is mainly found in the nucleolus and to a minor extent in the nucleoplasm (Suppl. Fig. 1C), corresponding to previous reports on TRAIP localization (15,33). Furthermore, a nucleolar and nuclear restricted biotin signal was observed using streptavidin-TexasRed while control cells expressing BirA*-HA showed only a signal outside of the nucleus (Suppl. Fig. 1D). These results demonstrated that TRAIP-BirA*-HA was functionally undistinguishable from TRAIP-HA.

In the next step, we purified biotinylated proteins from HeLa cells transfected either with TRAIP-BirA*-HA or BirA*-HA as described above. Two independent experiments were carried out and a representative example of isolated proteins detected with streptavidin-TexasRed and Coomassie Blue staining is shown in Suppl. Figs. 2A-2B. Purified biotinylated proteins were identified by LC-MS/MS analysis and validated with the Scaffold software. The results of comparing TRAIP-BirA*-HA vs. BirA*-HA from two independent experiments in asynchronous HeLa cells identified 385 proteins found in the TRAIP-BirA*-HA but not in the BirA* samples, whereby 229 (59.5 %) were identified in both experiments (Suppl. Fig. 2C), indicating that the two experiments were reproducible. We used the following criteria to establish the BioID dataset of highly significant hits enriched in the TRAIP-BirA* samples for further analysis: 95% protein threshold, 90% peptide threshold and minimum 2 peptides and p≤0.01 using Fisher’s Exact Test with Benjamini-Hochberg correction. The analysis produced a BioID dataset with 64 highly significant TRAIP interacting proteins. GO analysis revealed that 49 proteins localized to the nucleus and 33 to the nucleolus, in accordance with the previously reported TRAIP localization to the nucleus/nucleolus compartment. Inspection of the identified proteins revealed that a large proportion of proteins were implicated in DNA repair (Table 1). High confidence interactors such as PRKDC (DNA-PKcs), XRCC5 (Ku80) and XRCC6 (Ku70) and SMARCA5 (SNF2H) are preferentially implicated in the NHEJ branch of DSB repair (2). Importantly, in addition to DNA repair factors and DNA remodeling enzymes, ubiquitin E1 activating enzymes, E2 conjugating enzymes or ubiquitin were identified as putative interactors.

**Table 1.** Results of the BioID screen with TRAIP in HeLa cells. Total Spectrum counts are shown in the columns on the right. The following parameters have been used: 95% protein probability, 90% peptide threshold, 2 peptides minimum and the GO term ‘DNA repair’. Fisher’s exact test with Benjamini-Hochberg correction was used for statistical analysis (p-value).

| Identified Proteins | Uniprot | p-value | Experiment 1 |  | Experiment 2 |  |
| --- | --- | --- | --- | --- | --- | --- |
|  |  |  | TRAIP | BirA | TRAIP | BirA |
| DNA-dependent protein kinase catalytic subunit | P78527 | <0.0001 | 11 | 0 | 54 | 0 |
| X-ray repair cross-complementing protein 6 | P12956 | 0.00014 | 9 | 1 | 34 | 3 |
| X-ray repair cross-complementing protein 5 | P13010 | <0.0001 | 14 | 0 | 24 | 1 |
| Pre-mRNA splicing factor SYF1 | Q9HCS7 | <0.0001 | 11 | 0 | 13 | 0 |
| Sucrose non-fermenting protein homolog 2 | O60264 | 0.00031 | 7 | 0 | 14 | 0 |
| Transitional endoplasmatic reticulum ATPase | P55072 | 0.00062 | 7 | 2 | 35 | 3 |
| DNA polymerase delta catalytic subunit | P28340 | 0.0021 | 7 | 0 | 9 | 0 |
| Pachytene checkpoint protein 2 homolog | Q15645 | 0.0031 | 4 | 0 | 11 | 0 |
| DNA (-apurinic or apyrimidinic site) lyase | P27695 | 0.0045 | 9 | 0 | 5 | 0 |

### Ku80 and SMARCA5 are TRAIP interactors

Since the Bio-ID approach is a screening method (21), we set out to validate the interactors of TRAIP found in the BioID dataset (Table 1). Therefore, Ku80 and SMARCA5 were further investigated to see whether they bind to TRAIP. Co-immunoprecipitations with anti-FLAG M2 affinity gel using HeLa cells co-expressing HA-TRAIP and either EGFP-FLAG-Ku80 or SMARCA5-FLAG-Myc were performed. The results showed that TRAIP interacted with Ku80 (Fig. 1A) and SMARCA5 (Fig. 1D). Treatment with neocarzinostatin (NCS), a well-established radiomimetic compound causing preferentially DSBs (34), was used to investigate whether the interaction between TRAIP and Ku80 or SMARCA5 was affected by DNA damage in time course experiments. The TRAIP-Ku80 interaction was increased 30 min after NCS addition but was not significantly affected at later time-points (Figs. 1B-1C). In contrast, NCS significantly decreased the TRAIP-SMARCA5 interaction at all investigated time-points (30 to 480 min) (Figs. 1E-1F). These results indicate that the interaction of TRAIP with Ku80 or SMARCA5 is regulated by NCS.

**Fig. 1.**
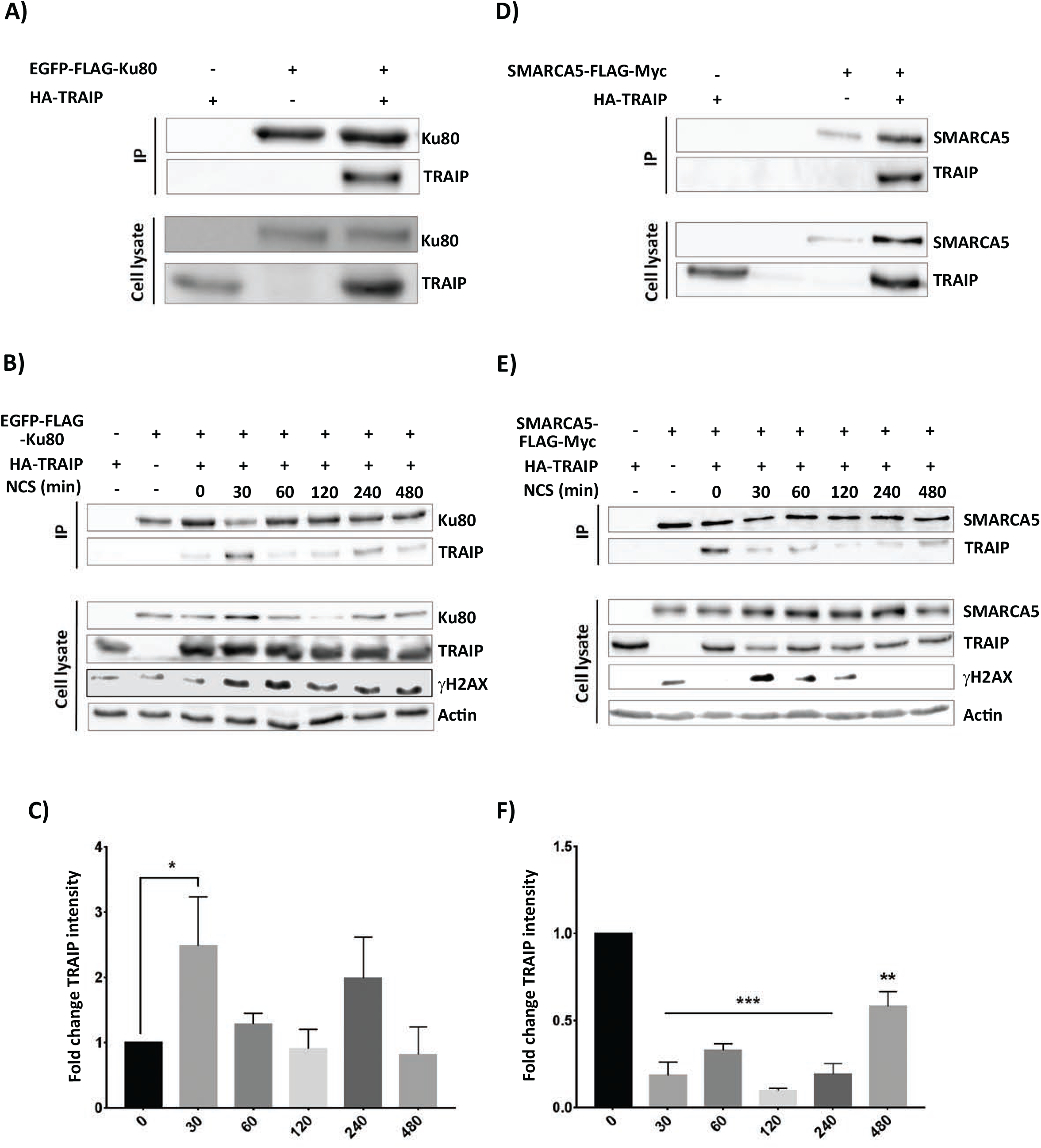
Identification of TRAIP interaction with Ku80 and SMARCA5. HeLa cells were transfected with HA-TRAIP and either EGFP-FLAG-Ku80 (**A, B**) or SMARCA5-FLAG-Myc (**D, E**). For the experiments shown in (**B, E**), cells were treated with NCS (200ng/ml) and cell extracts prepared at the indicated time-points. CoIPs were performed with anti-FLAG antibody and analyzed by immunoblot using anti-HA and anti-EGFP antibodies. Quantifications of TRAIP signal intensity of NCS-treated relative to the non-treated control (time 0 min) from the CoIPs with EGFP-FLAG-Ku80 (**B**) and SMARCA5-FLAG-MYC (**E**) are shown in (**C)** and **(F**), respectively. Results are depicted as mean±SD from 4 (**C**) or 2 (**F**) independent experiments. Statistical significance was calculated by two-sided Mann-Whitney test (**C**) and One-way ANOVA followed by Dunnett’s multiple comparisons test (**F**). Anti-γH2AX and anti-actin (**B, E**) were used to assess the NCS-dependent DNA damage and to ensure equal loading, respectively, using whole cell lysates.

### TRAIP is a constitutive DNA binding protein which localizes rapidly to sites of DNA damage

DNA repair factors accumulate to DNA damage sites after the induction of DNA damage (28,35). Therefore, we investigated whether TRAIP was detectable in the chromatin fraction after DNA damage. HeLa cells were fractionated into soluble proteins (S1), RNA bound proteins (S2) and chromatin (P2) (28). To ensure the purity of the different fractions, marker proteins such as α-tubulin (S1) and nucleolin (S1 and P2) were used (Fig. 2A). TRAIP was detected only in the fraction P2 revealing that it mainly binds to chromatin (Fig. 2A). At the global level, binding of TRAIP to chromatin was constitutive and independent of the induction of DNA damage by NCS (Fig. 2A).

**Fig. 2.**
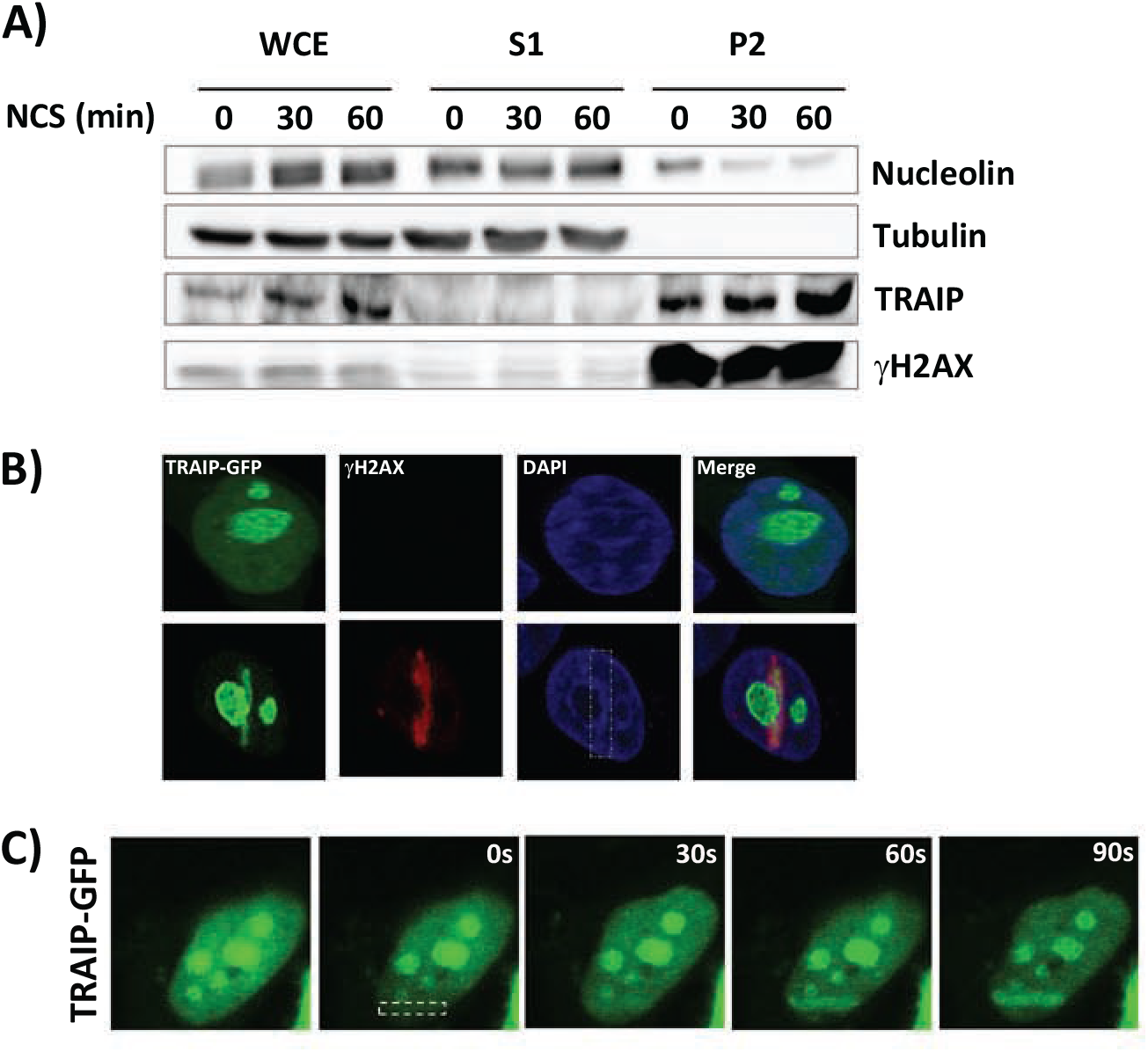
TRAIP-GFP localizes to sites of DNA damage. **A)** HeLa cells were treated or not with 200ng/ml NCS and collected at the indicated time points. Cells were lysed in denaturing buffer (WCE) or subjected to cell fractionation (S1 and P2) (28). Protein expression was analyzed with the indicated antibodies. **B)** TRAIP-GFP expressing HeLa cells treated (lower panel) or not (upper panel) with laser-microirradiation in the indicated area (dashed rectangle) were fixed 10 min later and stained for γH2AX and DNA (DAPI). **C)** Kinetic analysis of TRAIP-GFP accumulation at DNA lesions after laser-microirradiation (dashed rectangle) using live cell imaging. Time is indicated in seconds.

Furthermore, we examined whether TRAIP was specifically localized to DSBs after the induction of DNA damage. TRAIP-GFP transfected HeLa cells treated with NCS (200ng/ml) exhibited a partial overlap between γH2AX and TRAIP-GFP foci (data not shown). To answer this question more precisely, laser-microirradiation of BrdU-pretreated TRAIP-GFP transfected HeLa cells was used (36). TRAIP-GFP colocalized with γH2AX in the laser-irradiated area (Fig. 2B). Time-course experiments of TRAIP accrual at the DNA damage sites demonstrated that TRAIP was found within 1 min at laser microirradiation-induced DNA lesions (Fig. 2C). The rapid localization of TRAIP to DSBs after laser microirradiation suggests its implication in early events of DNA damage repair. These results are in line with previously published observations (17,19,20).

### TRAIP is important for activation of ATM

Based on the rapid recruitment of TRAIP in the laser-microirradiation experiments, we investigated the function of TRAIP during the early phase of DDR by analyzing the effect of TRAIP depletion on the DNA damage-mediated activation of ATM (4). HeLa cells depleted of TRAIP by lentiviral transduction with shRNAs (sh742 and sh743) were treated with NCS (200ng/ml), and ATM phosphorylation at Ser1981 was followed over time. After treatment with NCS, TRAIP depleted HeLa cells exhibited diminished levels of pS1981-ATM at all time points compared to control cells transduced with scrambled shRNA (SHC) (Fig. 3A). To further corroborate this data, we used fibroblast derived from a microcephalic primordial dwarfism patient harboring the homozygous hypomorphic TRAIP R18C mutation (17). Upon treatment of patient and normal fibroblasts with NCS (200ng/ml), strongly reduced ATM p1981 phosphorylation at 30 and 60 min post-treatment was observed in patient compared to normal fibroblasts (Fig. 3B). These data showed that TRAIP is crucial for the activation of ATM during the repair of NCS-induced DNA damage.

**Fig. 3.**
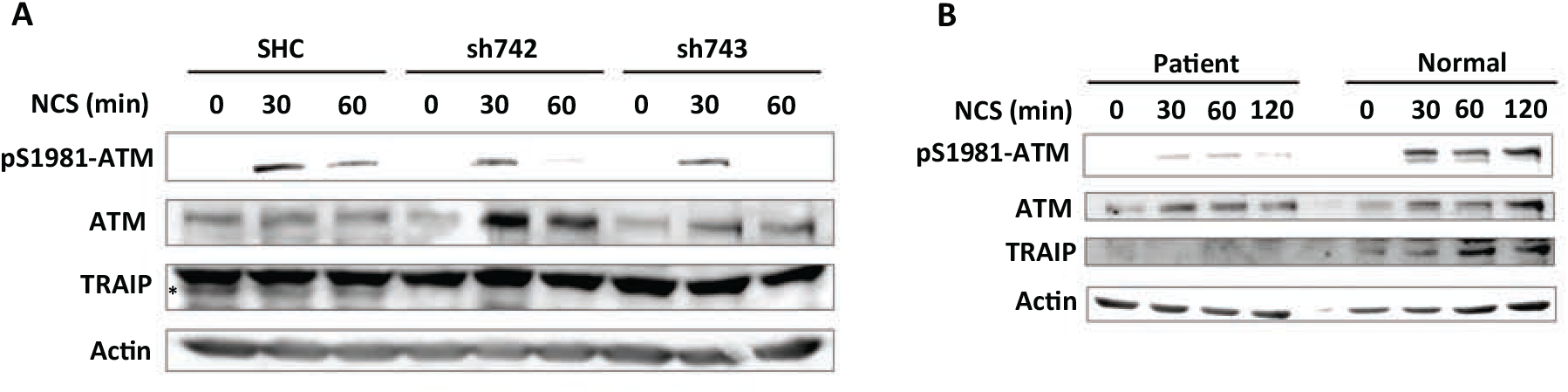
TRAIP is required for NCS-induced ATM activation. **A)** HeLa cells transduced either with control (SHC) or TRAIP-targeting (sh742, sh743) shRNAs were treated with NCS (200ng/ml). ATM-pS1981, ATM and TRAIP were detected at the indicated time-points by immunoblot. The asterisk denotes the TRAIP band. **B)** MPD patient (TRAIP R18C) and normal fibroblasts were treated with NCS (200ng/ml). ATM-pS1981, ATM and TRAIP were detected at the indicated time-points by immunoblot. Equal protein loading was assessed by anti-actin Western blots.

### TRAIP is implicated in crucial phosphorylation events during the DDR

We investigated whether reduced ATM kinase activity in TRAIP-depleted cells would lead to decreased phosphorylation of H2AX and Chk2, two downstream targets of ATM kinase activity (37). HeLa cells were transduced with control shRNA SHC or TRAIP targeting shRNAs sh742 or sh743. Analysis by qRT-PCR showed that TRAIP mRNA levels were reduced to 30% (sh742 and 17% (sh743) of the control. After treatment with NCS (200ng/ml), cells were fixed after different time points and quantified for mean γH2AX foci count and mean γH2AX foci intensity per nucleus by immunostaining (Figs. 4A and 4B). The γH2AX foci number per nucleus after one and two hours were significantly diminished by 34% and 45% (sh742) and 40% and 39% (sh743) in comparison to control cells, respectively (Fig. 4A). Similar results were obtained when the foci intensity per nucleus was quantified (Fig. 4B). Representative immunofluorescence images of γH2AX foci formation over time for control and TRAIP knock-down cells (sh743) are illustrated in Fig. 4C. In addition, immunoblot analysis of γH2AX in TRAIP-depleted cells (sh742) showed attenuated γH2AX signals at all time-points compared to control cells (Fig. 4D). These data demonstrated that TRAIP reproducibly affected the formation and maintenance of γH2AX foci, suggesting that TRAIP has an early function in the repair of NCS-induced DSBs.

**Fig. 4.**
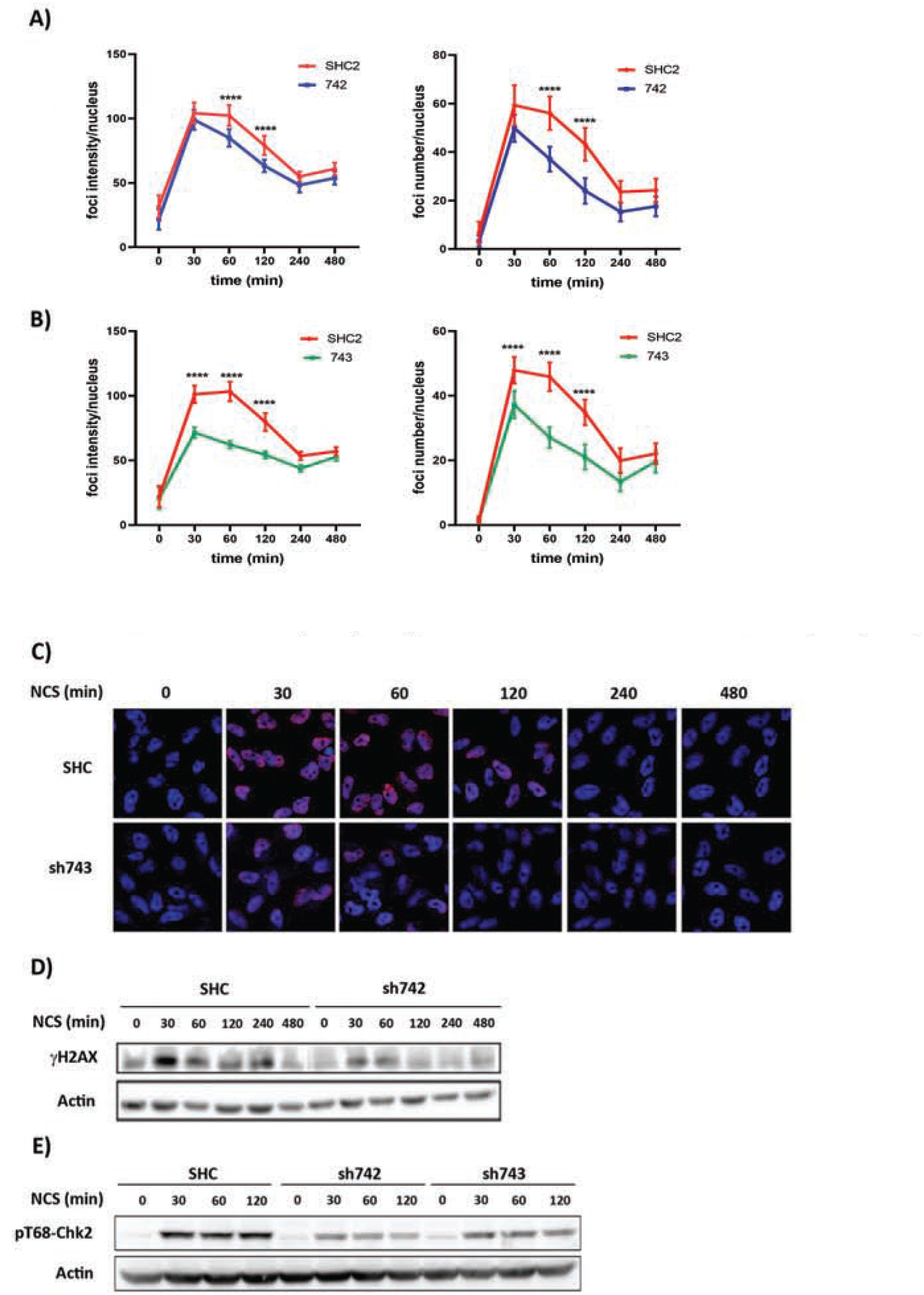
Knockdown of TRAIP in HeLa cells decreases γH2AX foci formation and Chk2 phosphorylation. Time course analysis (min) of the mean number of γH2AX foci/nucleus (**A**) or mean intensity of γH2AX foci/nucleus (**B**) after NCS (200ng/ml) treatment of HeLa cells transduced with control (SHC, gray dashed line) or TRAIP-targeting shRNAs (sh742 and sh743, black line). The results (mean±SD) for sh742 (left panels) and sh743 (right panels) are from 3 or 2 independent experiments with a total of n=210 or n=140 cells, respectively. Statistical significance was calculated by two-way repeated-measure ANOVA with Sidak’s correction; *p<0.5, **p<0.01, ***p<0.001, ****p<0.0001. **C)** Representative γH2AX immunofluorescence staining of TRAIP depleted (sh743) or control cells at the indicated time points (min) after treatment with 200ng/ml NCS. **D)** Immunoblot analysis of γH2AX after NCS (200ng/ml) treatment in TRAIP-depleted (sh742) and control (SHC) cells. **E)** HeLa cells were transduced with either control (SHC) or TRAIP targeting shRNAs (sh742 or sh743). Cells were treated with NCS (200ng/ml) 48 hours after transduction. The level of Chk2-pT68 was measured at the indicated time-points by immunoblot. A representative immunoblot from 4 independent experiments with similar results is depicted. Equal protein loading on immunoblots in (**D**) and (**E**) was assessed by actin staining.

DNA damage induced ATM kinase activity phosphorylates Chk2 at the residue T68 (37). Immunoblot analysis showed that NCS (200ng/ml) triggered Chk2 T68 phosphorylation within 30 min and the signal persisted for at least 120 min (Fig. 4E). In contrast, TRAIP knockdown in HeLa cells using either sh742 or sh743 resulted in decreased Chk2 T68 phosphorylation at all investigated time points (Fig. 4E). These results suggest that TRAIP is controlling not only the DNA damage response but also ATM-Chk2 signaling and therefore G2/M checkpoint activation.

### Correct DNA end use during NHEJ mediated repair depends on TRAIP

DSBs are repaired either by high fidelity homologous recombination (HR) or error-prone non-homologous end-joining (NHEJ). To study the role of TRAIP in HR and NHEJ, DSB-repair reporter assays using stable 293T cell lines with chromosomally integrated copies of the plasmids pDR-GFP (HR) or pimEJ5GFP (NHEJ) were performed (25,38). Stable 293T cells transduced either with TRAIP targeting (sh742, sh743) or control (SHC) shRNAs were cotransfected with pCBASce-I encoding I-SceI and pLVX-mCherry-N1 to assess transfection efficiency. The results showed that the relative number of GFP-positive cells was increased in pEJ5-GFP cells treated with sh742 and sh743 compared to control (Figs. 5A, and 5B left panel). As reported previously (38), inhibition of DNA-PKcs with NU7741 in pimEJ5GFP cells increased the number of GFP-positive cells (data not shown). In contrast, no significant differences in HR repair frequencies between control and TRAIP-depleted cells were observed (Figs. 5C, and 5D left panel). Immunoblot analyses confirmed strong reduction of TRAIP expression in cells transduced with either sh742 or sh743 (Figs. 5B and 5D, right panels). These results indicate that TRAIP is important to suppress distal end-joining but is not involved in HR-mediated repair.

**Fig. 5.**
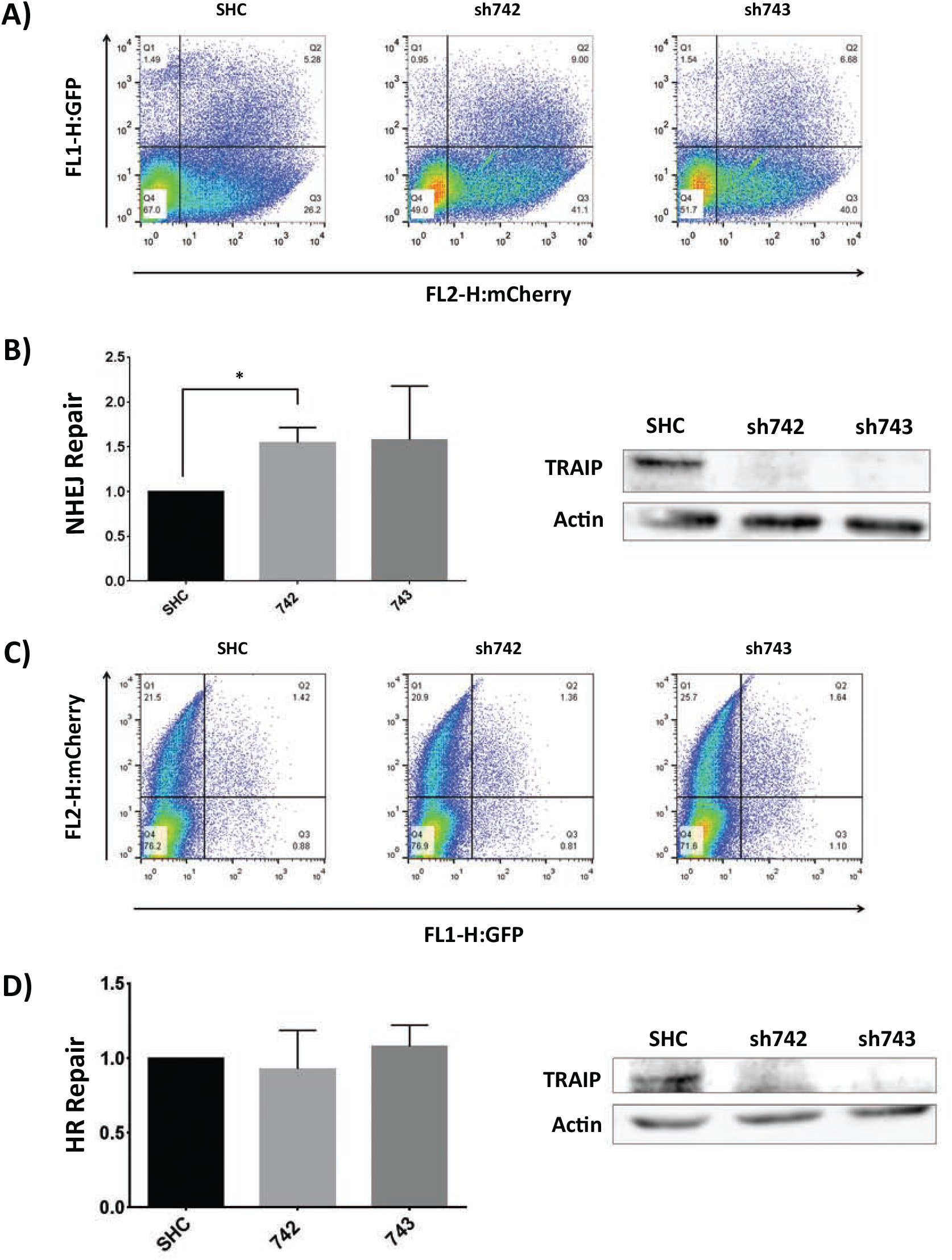
TRAIP is involved in NHEJ. **A)** Flow cytometry results of NHEJ DNA repair assay in stable pimEJ5GFP 293T cells depicted as scatter of plot GFP and mCherry positive cells. **B)** NHEJ repair efficiency shown as mean±SD from 3 independent experiments (left panel), and verification of TRAIP knockdown efficiency (right panel). Statistical significance was calculated with the Kruskal-Wallis test followed by Dunn’s multiple comparisons. **C)** Flow cytometry results of HR DNA repair assay in stable HR-GFP 293T cells depicted as scatter plot of GFP and mCherry positive cells. **D**) HR repair efficiency shown as mean±SD from 4 independent experiments (left panel), and verification of TRAIP knockdown efficiency (right panel).

## DISCUSSION

Microcephalic primordial dwarfism patients with homozygous hypomorphic TRAIP mutations show increased UV-induced DNA damage and impaired cellular proliferation (17), consistent with TRAIP knock-down experiments (14) and Traip KO animals (13). TRAIP interacts with PCNA to protect genome stability during replication-stress (18,19) and with RAP80 to ensure the recruitment of BRCA1 to DNA damage lesions (20). Furthermore, TRAIP regulates the SAC to maintain genome integrity (15,16). However, no molecular mechanisms how exactly TRAIP regulates DNA damage repair and the SAC, or bona fide substrates have been reported, encouraging the search for TRAIP interacting proteins.

The TRAIP Bio-ID screening identified proteins implicated in the DNA damage repair, chromatin remodeling, transcription and ubiquitylation. The identification of Ku80 and SMARCA5 as TRAIP-interacting proteins suggested a putative role of TRAIP in the NHEJ branch of DSB repair and/or chromatin remodeling. Interestingly, NCS treatment increased binding of TRAIP to Ku80 while binding to SMARCA5 was strongly diminished during 8 hours. The transiently increased interaction between TRAIP and Ku80 showing a maximum at 30 min after NCS addition might indicate that the interaction occurs at DSBs. Ku80 has to be removed rapidly after repair of DSBs to allow transcription and replication to proceed normally, which constitutes a topological problem because Ku80 encircles the DNA in a toroidal fashion (39). Ku80 has been shown to be ubiquitylated and neddylated in response to DSBs (40–42). The E3 ubiquitin ligase RNF8 has been implicated in K48-specific ubiquitylation of Ku80 (41), but recently no effect of RNF8 was found (40). It remains to be investigated whether TRAIP ubiquitylates Ku80 in response to DSBs. Recently, the AAA-ATPase VCP/p97, which has been found in the TRAIP BioID dataset (Table 1), has been implicated in the removal of K48-ubiquitylated Ku80 from DNA after DSB repair (43).

TRAIP was reported to bind to PCNA through a C-terminal PIP sequence (18,19) and to function as PCNA clamp unloader during DNA replication. In our BioID screen PCNA was not found as significant hit, which can most likely be explained by the BirA* addition at the C-terminal end of TRAIP, presumably blocking PCNA access to the PIP site of TRAIP (18,19).

Several DNA repair proteins show increased chromatin binding after induction of DNA damage (28). Cell fractionation experiments in NCS-treated asynchronous HeLa cells revealed that endogenous TRAIP associates constitutively to chromatin (Fig. 2A), consistent with previous immunofluorescence results from mitotic chromosomes (15), but can be rapidly mobilized to DNA damage sites. A recent report showed that TRAIP associates with chromatin to a similar extent in non- and camptothecin-treated cells (19). Laser microirradiation experiments with HeLa cells expressing TRAIP-GFP showed accumulation of TRAIP to the irradiated area within minutes (Fig. 2), in accordance with recent reports (17,19,20). Ku70/80 are recruited within seconds to DSBs (44), while TRAIP accumulates later suggesting that it acts downstream of Ku70/80. Interestingly, DNA repair factors such as 53BP1 and BRCA1, determining the DNA repair pathway choice, are recruited to DSBs with a similar timeframe as TRAIP-GFP (35).

ATM is a crucial protein for the initiation of chromatin remodeling processes, DNA repair and the activation of cell cycle arrest (37). TRAIP-depleted HeLa cells and fibroblasts from a MPD patient (homozygous hypomorphic TRAIP R18C mutation (17)) showed decreased ATM phosphorylation after NCS treatment (Fig. 3A and 3B). Previously, it has been reported that in TRAIP KD cells and MPD fibroblasts ATM activation was not impaired 4 and 24 hours after UV-C treatment compared to control cells (17). It remains to be seen whether the role of TRAIP in ATM activation depends on the DNA damaging agent. TRAIP seems to regulate ATM activation rapidly after induction of DSBs, consistent with the rapid accrual to laser-microirradiation lesions (Fig. 2C). These results strongly support the notion that TRAIP is necessary for the rapid activation of ATM during the repair of DSBs.

To investigate the consequences of the reduced ATM activation, we studied two well-known targets of its kinase function, H2AX and Chk2. Analysis of γH2AX revealed significant lower foci numbers and foci intensities in NCS-treated cells lacking TRAIP compared to control cells. Consistent with the time course of NCS-dependent ATM phosphorylation in TRAIP-depleted cells, the differences in the γH2AX levels were much larger at 1 and 2 hours than at 30 min after NCS addition, indicating that TRAIP may also affect the maintenance of γH2AX. These findings correspond to recent reports revealing decreased γH2AX signal in TRAIP-depleted cells undergoing replication stress (17,19). In contrast, slower disappearance of γH2AX foci was observed in TRAIP-depleted HeLa cells after ionizing radiation (20). Explanations for these contradictory observations are not known at the moment but needs further investigations.

TRAIP knock-down reduced NCS-dependent Chk2 T68 phosphorylation in HeLa cells (Fig. 4E) implying a function of TRAIP to establish the G2/M checkpoint. Our data regarding ATM/Chk2 signaling is consistent with a recent report using ionizing radiation (20). TRAIP regulates the spindle assembly checkpoint (SAC) by controlling Mad2 abundance at kinetochores/centromeres (15,16), localizes to centromeres (16) and prevents chromosome aberrations and aneuploidy (15,16). Interestingly, it has been reported that ATM-depleted cells proceed faster through mitosis, and chemical inhibition of ATM or knock-down of the ATM substrate MDC1 impaired Mad2 localization at kinetochores (45). Furthermore, Chk2 controls mitotic progression time and mitotic exit, is required for proper spindle formation and proper chromosome alignment and segregation (46,47). Mitotic kinases provide a negative feed-back system to modulate the ATM-Chk2 arm of DNA damage signaling (48), underlining the significance of crosstalk between DDR and SAC to preserve the genetic integrity (49). In summary, our results demonstrate that TRAIP regulates the DSB-dependent activation of ATM and downstream targets involved in DNA repair and the G2/M cell cycle checkpoint.

DNA repair assays measuring distal end-joining (NHEJ) and HR showed that TRAIP depletion increased distal end-joining but did not alter HR activity compared to control cells, implying that TRAIP inhibits distal end-joining. Similarly, inhibitors of ATM and DNA-PKcs increased also distal end-joining in stable EJ5-GFP U2OS cells (38). DNA-PKcs phosphorylates ATM to inhibit its kinase activity in response to DNA damage (50) whereas phosphorylation of DNA-PKcs by ATM is required for NHEJ repair of DSBs (9), indicating DNA-PKcs and ATM act cooperatively during DNA repair. The discovery that TRAIP depletion affects NHEJ repair and ATM kinase activity is consistent with TRAIP BioID data, which identified three core NHEJ components (Ku70, Ku80, DNA-PKcs). These results indicate that TRAIP may regulate the crosstalk between DNA-PKcs/ATM to control the choice of repair pathways. In addition, TRAIP may affect chromatin remodeling by SMARCA5, an ATP-dependent helicase which works with ATM to stabilize γH2AX in response to DSBs (51) and interacts with Ku80 in a DSB-dependent manner (52). Since ATM signaling is implicated in HR repair (53), it was surprising that TRAIP depletion had no effect on HR, contrary to what was reported previously in U2OS cells (20). The residual ATM activity in TRAIP-depleted 293T cells may be sufficient to perform HR but not NHEJ because Ku80- or SMARCA5-dependent steps of NHEJ could be affected as well.

In summary, we reveal novel TRAIP interactors (Ku80 and SMARCA5) and TRAIP’s role in regulating ATM activation crucial to efficiently repair DSBs, which is consistent with the negative effect of reduced TRAIP expression on genomic integrity (15,16).

## ACCESSION NUMBERS

Mass spectrometry data has been deposited to the ProteomeXchange repository with the project accession number PXD006715 (project DOI 10.6019/PXD006715).

## SUPPLEMENTARY DATA

Supplementary Data are available at NAR online.

## ACKNOWLEDGEMENTS

We thank all the investigators mentioned in Material and Methods for their gifts of plasmids.

---

**Suppl. Fig. 1. Functionality and correct localization of TRAIP-BirA*-HA. A)** Cell extracts from FLAG-ubiquitin, TRAIP-BirA*-HA or HA-TRAIP transfected 293T cells were immunoprecipitated with anti-HA antibody. Immunoblot analysis by were carried out using anti-HA or anti-FLAG antibodies. **B)** Immunoblot with streptavidin-HRP or anti-HA antibodies in TRAIP-BirA*-HA and BirA*-HA transfected cells cultured for 24 hours with or without 50μM biotin. **C)** Confocal immunofluorescence showing nucleolar localization of TRAIP-BirA*-HA transfected HeLa cells detected with anti-HA or anti-TRAIP antibodies. **D)** Confocal immunofluorescence using anti HA-antibodies or streptavidin-TexasRed (TR) in HeLa cells transfected with TRAIP-BirA*-HA (upper panels) or BirA*-HA (lower panels) with or without pre-extraction. Cells were preextracted with 0.2% Triton-X100 for 20 sec before fixation and DNA was stained with DAPI (C and D).

**Suppl. Fig. 2. Pull-down and mass-spectrometry analysis of biotinylated proteins. A)** Immunoblot of biotinylated proteins purified from HeLa cells transfected with TRAIP-BirA*-HA or BirA*-HA using streptavidin-HRP. **B)** Colloidal Coomassie staining of the purified proteins shown in A. **C)** Venn diagram of TRAIP BioID and the BirA* BioID proteins identified in experiments 1 and 2. The BirA* BioID data from experiments 1 and 2 were collated.

