## Supplementary figures and images for "TRAIP regulates DNA double-strand break-induced ATM activation"

### Supplemental Figure 1

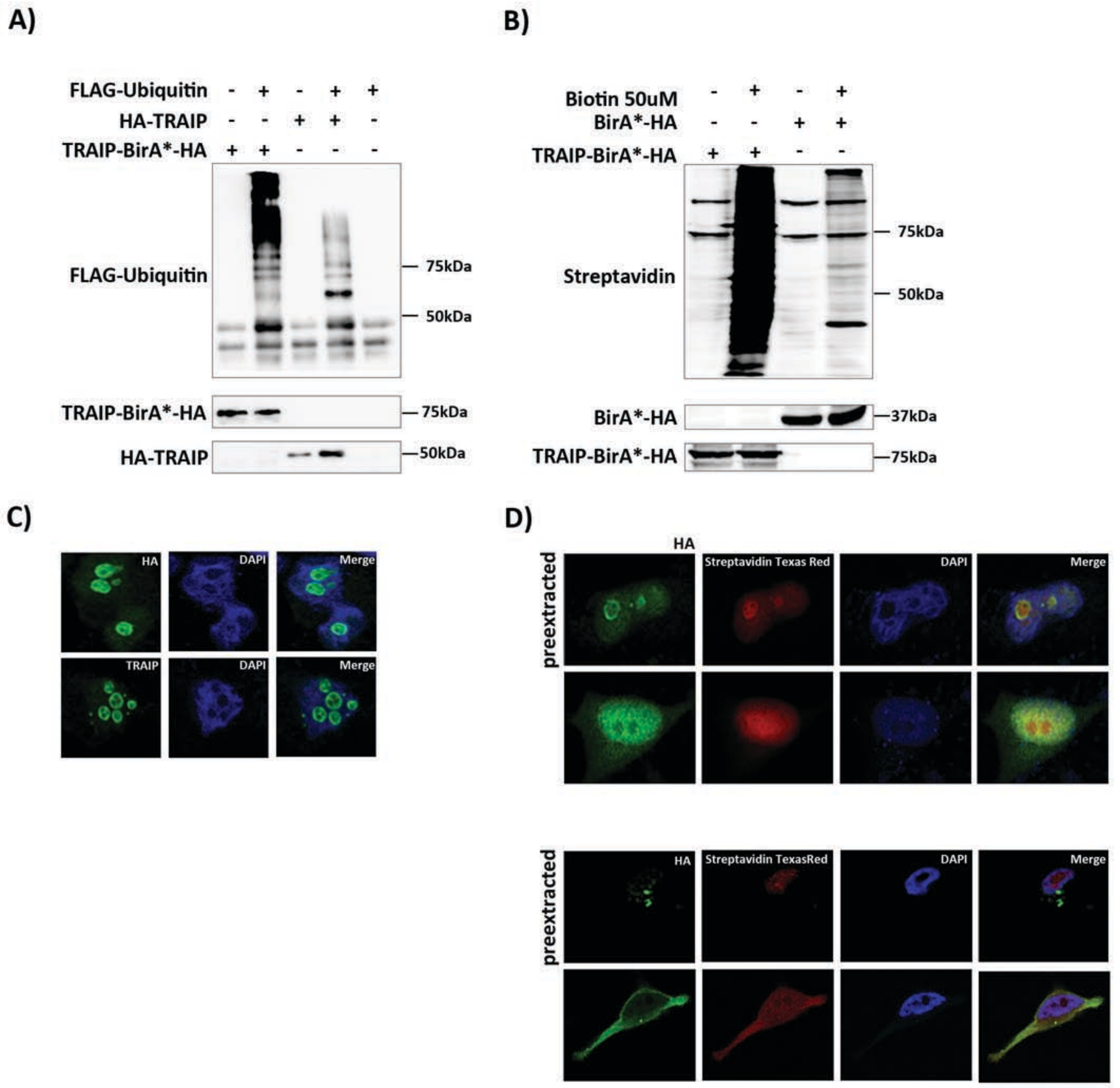

### Supplemental Figure 2

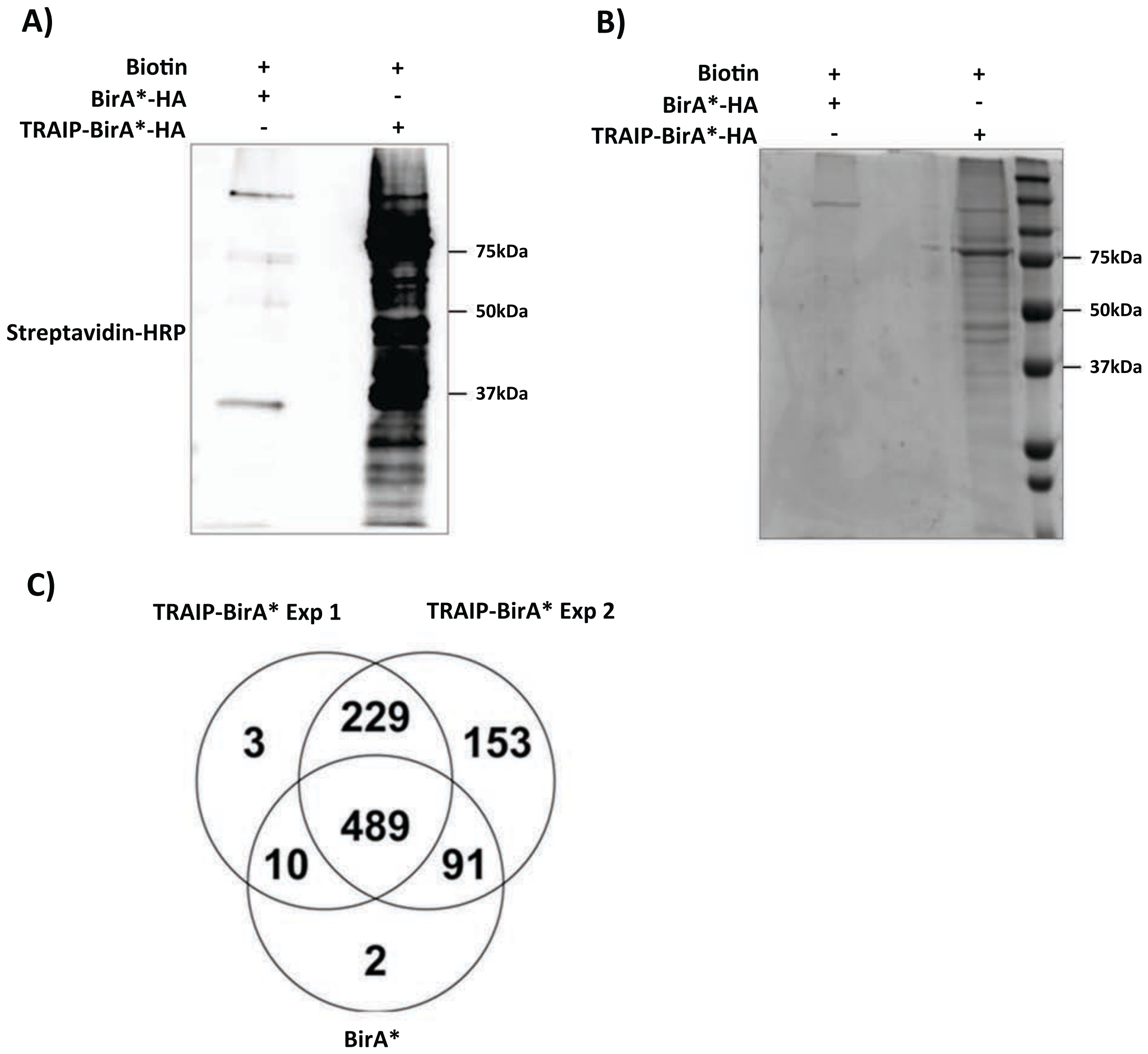
